# Pindel-TD: a tandem duplication detector based on a pattern growth approach

**DOI:** 10.1101/2023.10.08.561441

**Authors:** Xiaofei Yang, Gaoyang Zheng, Peng Jia, Songbo Wang, Kai Ye

## Abstract

**Tandem duplication** (TD) is a major type of **structural variation** (SV), and plays an important role in novel gene formation and human diseases. However, TDs are often missed or incorrectly classified as insertions by most of modern SV detection methods due to the lacking of specialized operation on TD related mutational signals. Herein, we developed a TD detection module of Pindel referred as Pindel-TD based on a TD specific **pattern growth** approach. Pindel-TD detects TDs with a wide size range at single nucleotide resolution. Using simulation and real read data of HG002, we demonstrate that Pindel-TD outperformed other leading methods in terms of precision, recall, F1-score and robustness. Further applying Pindel-TD on data generated from K562 cancer cell line, we identified a TD located at the seventh exon of *SAGE1*, explaining its high expression. Pindel-TD is available at https://github.com/xjtu-omics/pindel and free for non-commercial use.

## Introduction

Tandem duplication (TD) is one of the major types of structural variation (SV) [1, 2], contributing to *de novo* gene structure formation [3], evolution of biosynthetic pathway in plants [4], and human diseases, including autism [5] and cancers. Cancers, such as breast, ovarian, and endometrial carcinomas, can be further divided into different tandem duplicator phenotype (TDP) subgroups based on the frequency and distribution of TDs [6, 7]. TDs contribute to tumorgenesis by augmenting oncogene expression and disrupting tumor suppressor genes [6, 7].

Fueled up by the development of the next and third-generation sequencing technology, a lot of SV detection methods have been developed based on model-matched or model-free strategies. In general, methods follow a model-matched strategies by first extracting the mutational signal, such as discordant read-pairs, read-depth, and split-reads, and then matching signal to a specific SV model. For example, Pindel applied a pattern growth approach to take advantage of the split-read signal to detect SVs with precise breakpoints [8, 9]. Delly performed an integration analysis of the pair-end mapping and split-reads information to detect SVs at single-nucleotide resolution [10]. Lumpy was developed to precisely identify SVs based on a general probabilistic framework to integrate three different mutational signals of read-pair, split-read, and read-depth [11]. Model-matched SV detection strategy has been proved to be efficient on detecting simple SVs, *e*.*g*. insertion, deletion, and inversion, but lacks of performance on complex SVs (CSVs), which contain multiple breakpoints and playing important roles in cancer [9, 12]. Mako [13] and SVision [14] are two methods designed based on model-free strategy to detect CSVs from the short-read sequencing data and long-read sequencing data, respectively. Although these methods perform well on detecting the most frequent types of SVs, like insertions and deletions, certain type of SVs, like TDs, are detected with lower accuracy and lower recall rate, *e*.*g*. researchers have pointed out that TDs are frequently reported as insertions [15, 16], due to the lacking of specific optimization on TDs. Therefore, developing a TD detection tool by specifically optimizing to accurate characterize TDs is an eminent need in the genomic community.

Here, we reported a TD detection module of Pindel method referred as Pindel-TD, in which we specifically optimized the pattern growth approach applied in Pindel to satisfy the short-read alignment signatures of TDs with a wide size range. We also applied split-read analysis in Pindel-TD to acquire single nucleotide resolution of the breakpoint. We evaluated the performance of Pindel-TD on both simulated sequencing data and real sequencing data of sample HG002 [17], demonstrating that Pindel-TD is effective for TD detection and outperforms other methods, *e*.*g*. Manta [18], Delly [10], Lumpy [11], and DINTD [19]. We also reported the results of Pindel-TD using short-read sequencing data of a cancer cell line K562 from ENCODE [20] to illustrate its potential application on cancer genomic data.

## Methods

### Overview of Pindel-TD

We proposed a pattern growth approach for TD detection with different sizes and implemented Pindel-TD in C++. Overall, there are four steps for TD detection(**Figure 1A**), following the framework of pattern growth strategy in Pindel for the detection of large deletion and small insertion [8]. Firstly, we selected the read-pairs with only one read mapped uniquely (mapped only with ‘M’ character in its CIGAR string) while its mate showing split-read (soft clipped). For each selected read-pair, the mapped read with a high mapping quality (larger than the *anchor_quality* parameter, default is 30) was considered as a reliable anchor read (*e*.*g*. blue arrows in **Figure 1**), determining the searching direction of subsequent split read analysis of soft clipped read. Secondly, we apply a pattern growth approach to find minimum (the *min_distance_to_the_end* parameter in Pindel-TD, default is 8) and maximum unique substring start from either the leftmost or the rightmost of the unmapped read (showing soft clip in BAM, *e*.*g*. black arrows in **Figure 1**). Next, we carefully processed the split-read information to identify the TDs with accurate breakpoints. Finally, we removed the redundant TDs according to their length and break points to get final TD set.

**Figure 1.**
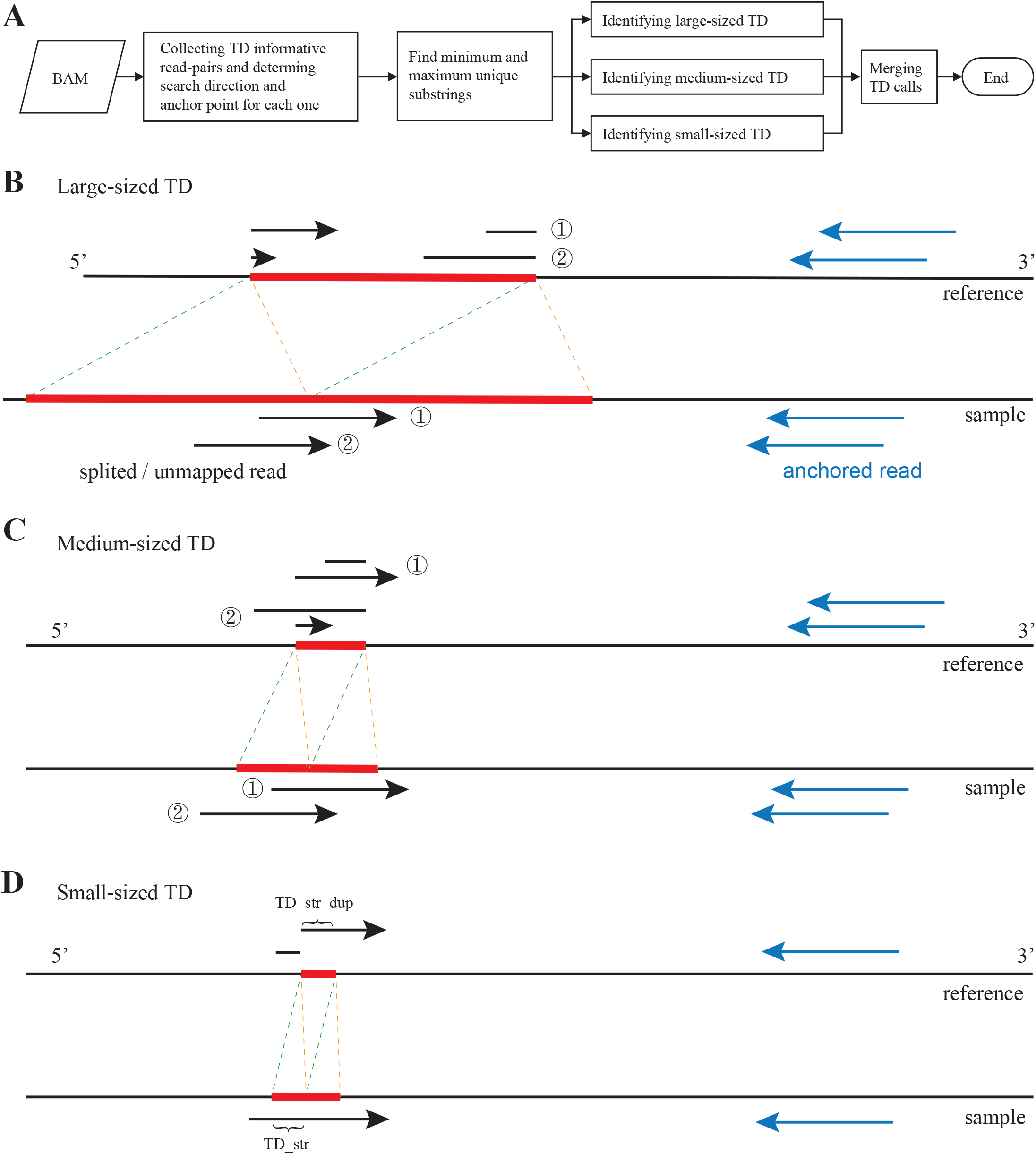
Pindel-TD framework. **A**. The workflow of Pindel-TD. **B-D**: The Characterization of the alignment signals of large (B), medium (C) and small (D) TDs. The red blocks indicated the TD regions in reference and sample genome green and orange dash lines indicated the TD boundaries blue and black arrows indicated the anchor reads and the unmapped reads of the TD informative read-pairs, respectively. Two segments of the unmapped read for each TD informative read-pair on reference indicated the detected minimum and maximum substrings using to determine the breakpoint. In (D), TD_str and TD_str_dup indicated the TD sequences inferred from the unmapped read.

### Detail steps for TD detection

We defined large, small, and medium-sized TDs as following: large-size: *len* (*TD*) > *read* _ *len* medium-size: *read* _ *len* > *len* (*TD*) > 1/2 × *read* _ *len* ; small-size: *len* (*TD*) < 1/2 × *read* _ *len*, where *read_len* indicates the read length, *len(TD)* indicates the length of a TD event. We designed four steps to detected TDs under a wide-size spectrum (**Figure 1**).

1. For each TD informative read-pair, we defined the anchor point as the 3′ end of the mapped read
2. Applying pattern growth approach to search for minimum and maximum unique substrings from the 3′ end of the unmapped read within the range of two times of insert size from the anchor point.
3. Applying pattern growth approach to search for minimum and maximum unique substrings from the 5′ end of unmapped read within a region. For large-size TDs, the region is defined as a range of *Max_TD_Size* (a user defined parameter) from the already mapped 5′ end of the unmapped read obtained in step 2. For medium and small-size TDs, the region is defined as the range of 2 times of *Max_Span_Size* (a user defined parameter) with the center being the already mapped 5’ end obtained from step 2. For small-size TDs, we stopped searching once the current searching coordinate is the same as the coordinate of the already mapped 5’ end obtained from step 2.
4. For large and medium-size TDs, combining of substrings extracted from step 2 and step 3 and checking whether a complete unmapped read can be reconstructed. If yes, Pindel-TD records two substring ends as breakpoints of a large-size TD and stores the interval into the candidate TD database. For small-size TDs, Pindel-TD extracts the candidate duplication sequence by cutting the same length sequence from the maximum unique substring obtained in step 2 at 5′ end (TD_str_dup in **Figure 1D**). To get reliable small-size TDs, Pindel-TD calculates the edit distance between TD_str and TD_str_dup by Edlib (https://github.com/efcs/elib) [21]. If the edit distance is smaller than 4, Pindel-TD stores it in the database.

All records in the candidate TD database are sorted by the 5′ reference coordinate. Candidate TDs with at least two supported read is considered as reliable TDs.

### Merging TD calls

Since the alignment signals of the above three strategies may be overlapping, it is necessary to merge TD calls. For a TD call *TD*_*i*_, we identified its overlapping TD callset by checking whether the range of a TD call (*TD*_*j*_) overlapping with *TD*_*i*_ and |*len* (*TD*_*i*_) − *len* (*TD*_*j*_) | < 2 × min (*len* (*TD*_*i*_), *len* (*TD*_*j*_)), where *len* (*TD*_*i*_) is the length of *TD*_*i*_. For an identified set, we selected the most frequent TD call as the represented TD event and reported out. This process was executed iteratively to get the final TDs.

### Simulated data generation

In order to evaluate the performance of Pindel-TD, we first simulated different sizes of TDs on human chromosome 1 by VISOR v1.1 [22]. We first apply “*randomregion*.*r*” script in VISOR to generate 37 BED files on human chromosome 1 (Table S1). Each BED file contains 100 simulated TDs with a specific length. The simulated TD length varies from 10 bp to 9 Kb (Table S1). Then “*VISOR HACk -b $TD_BED -g human_chr1*.*fa -o $OUTPUT*” command was used to put the simulated TDs into human chromosome 1 of GRCh38. Next, we employed wgsim version 1.12 (https://github.com/lh3/wgsim) to simulate 15x coverage of paired-end sequencing data of the human chromosome 1 containing simulated TDs with 100 bp read length and 400 bp insert size. In the simulated sequencing, we set the error rate as 0.5%. We aligned the simulated sequencing data to GRCh38 chromosome 1 by BWA-MEM v0.7.17-r1188 [23] to get the BAM file for TD calling. We applied *bedtools intersect* to check whether a simulated TD is recovered by different methods.

### Processing the benchmark SV set of real data (HG002/NA24385)

The released benchmark SV set (version v0.6) of HG002/NA24385 [17] was constructed by integration of SV callsets from 19 SV detection methods and four sequencing technologies, including short-read, linked-read, long-read, and Bionano optical map, against GRCh37 [17]. In this benchmark, the SVTYPE only labels INS (insertion) and DEL (deletion), and the duplication (both tandem and dispersed duplication) is reported as DUP in REPTYPE. In total, there are 2,817 duplications, and 1,882 of which are labeled as Illcalls (calling from Illumina short-read sequencing data).

To identify TDs from the duplications, we manually inspected the dotplots created by Gepard version 2.1.0 [24] using the DUP related long-read extracted from the BAM file generated by ngmlr [25] with parameter “*-x pacbio*” based on the High-Fidelity reads and the related GRCh37 sequences. Furthermore, we excluded the candidate TDs located at repetitive regions of GRCh37 since detection of TDs in these regions is still challenging for short-read based methods [1]. Eventually we obtained 600 non-repetitive benchmarked TDs (Table S2). We applied Truvari *bench* subcommand with parameters “*-r 1000 -p 0*.*0 -P 0*.*5 -s 50 -S 50 --sizemax 2000 --typeignore*” (v3.5.0) [26] to compare the TD calls reported by various tools with the benchmark.

## Results

### Performance evaluation on simulated data

To evaluate the performance of Pindel-TD on TD detection, we examined how well it can recover the simulated TDs and compared the results with that from Manta, Delly, and Lumpy. We calculated the precision, recall and F1-score of each TD set with different sizes (Table S3). We found that Pindel-TD achieved the best performance on simulated TDs with average precision, recall and F1-score of 97%, 94%, and 96%, respectively, while Manta, Lumpy, Delly, and DINTD ranked the second, third, fourth and fifth with the average F1-score of 92%, 92%, 68%, and 31%, respectively (**Figure 2A** and Table S3). The average false discovery rate of Pindel-TD is about 3% (Table S3). The most significant improvement of Pindel-TD compared to other methods was the detection of TDs with length smaller than read length (100 bp) (Table S3). In addition, Pindel-TD extended the lower bound of the detectable TDs from 30 bp (Manta) to 10 bp. For TDs smaller than 30 bp, Pindel-TD achieved average F1-score of 92% (Table S3). Delly and Lumpy lacks the ability for detecting small-sized TDs, *e*.*g*. Delly missed all TDs smaller than 60 bp and Lumpy missed all TDs smaller than 100 bp. For TDs larger than 500 bp, we found that Manta obtained slightly higher average F1-score than Pindel-TD (97% for Manta and 95% for Pindel-TD), might due to the probability of finding a unique substring decreases when we enlarge the parameter *Max_TD_Size* in step 3. All the simulation evaluation results indicated that Pindel-TD is an effective and robust method for detecting TDs given the whole size range.

**Figure 2.**
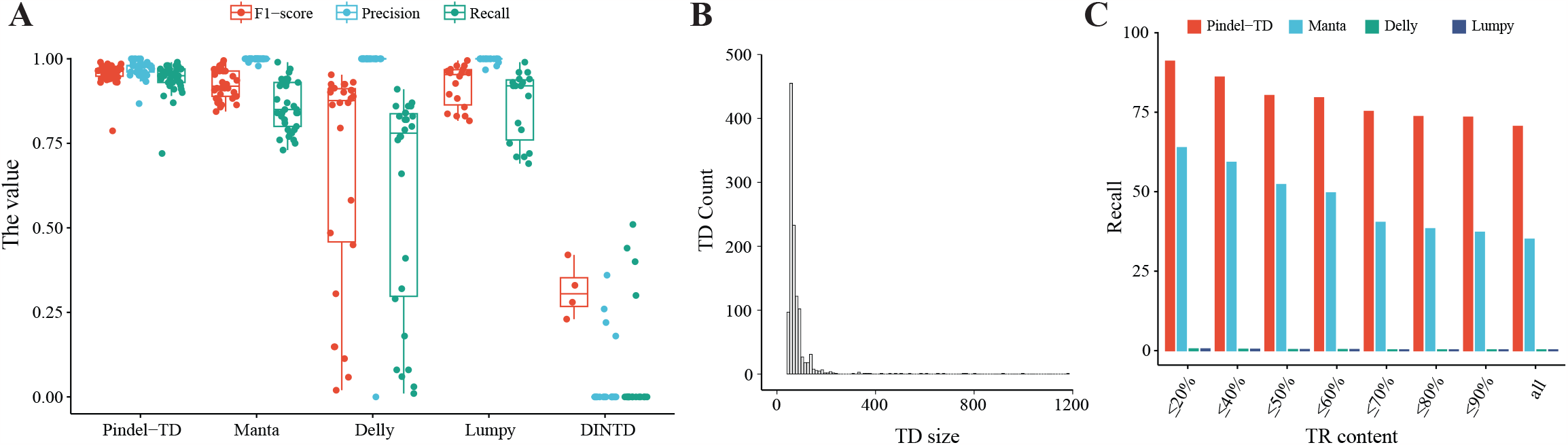
The performance evaluation on simulation and real data. **A**. The F1-score, precision, and recall on the simulated data for four TD detection methods. The detail values are shown in Table S1. **B**. The length distribution of 1,162 TDs detected from HG002 data. **C**. The recall of four methods on HG002 benchmarked TDs for different TR content. TD: tandem duplication, TR: tandem repeat.

### Performance evaluation on HG002 data

We further examined the performance of Pindel-TD on paired-end sequencing data of the well-studied cell line HG002/GM24385. We applied Pindel-TD on the short-read sequencing data of HG002 and detected 1,162 TDs with length larger than 50 bp (**Figure 2B**, Table S4). We found that 955 (82.19%) were overlapped with the released benchmark SV set (v0.6), higher than that from Manta (632 of 799, 79.1%), Delly (390 of 1032, 37.8%), and Lumpy (109 of 227, 48.2%) (Table S5). Moreover, we found that 70.5%, 35.0%, 0.2%, and 0.2% benchmarked TDs were recalled by Pindel-TD, Manta, Delly, and Lumpy, respectively (**Figure 2C**, Table S6), indicating Pindel-TD is effective on TD detection. Reference tandem repeat (TR) sequence introduces the TD-like alignment signals, affecting the accuracy of TD prediction (Table S6). We next investigated the influence of TR content on the detection of TDs for these four methods. We computed the TR content of each benchmarked TD by tandem repeat finder (TRF) [27], and calculated the recall for different TR content. We found the recall of Pindel-TD varied from 91.0% to 73.4% with TR content varying from ≤ 20% to ≤ 90%, while that of, the second best, Manta varied from 63.8% to 37.2%% (**Figure 2C**, Table S6), indicating that Pindel-TD is more robust than other methods for different TR content.

### Detecting TDs on K562 cell line

To test the potential usage of Pindel-TD on cancer genomic sequencing data, we applied it on the short-read sequencing data of a human erythroleukemic cell line K562 from ENCODE [20]. In total, we detected 888 TDs on autosomes and chromosome X, and 531 out of them were Pindel-TD specific TDs (Table S7). Some of the detected TDs related with protein-coding genes, *e*.*g. FLT3, MUC3A, SAGE1, PLIN4, CBS*, and *ACSF3*. We found that one TD (chrX:135,906,203-135,906,590) occurred on the seventh exon of *SAGE1*, which is a gene coding a cancer antigen (**Figure 3A**). We found that the expression of *SAGE1* in K562 cell line is significantly higher than that in T cell (*p*-value = 0.0036, Wilcoxon Rank Sum and Signed Rank test, **Figure 3B**, Table S8), especially the seventh exon acquired the most read coverage among its 20 exons (Figure S1), suggesting the TD directly increased the gene expression of *SAGE1* in K562 cell line. The tumor-specific expression pattern of *SAGE1* has been found in several solid cancers, *e*.*g*. bladder cancer, lung cancer, head and neck cancers, suggesting its role as a potential target for cancer immunotherapy [28]. A few studies have also reported the relevance between *SAGE1* and leukemia [29, 30], while the genetic variation of *SAGE1* has not been well studied in cancers. One of the Pindel-TD specific TD (chr21:43,063,062-43,063,130) affects the coding sequence of protein-coding gene *CBS*, which encodes cystathionine *β*-synthase. Higher expression level of *CBS* in chronic myeloid leukemia (CML) patients than that in control samples has been reported [31], while the underline genetic mechanism is not well studied. Our finding may provide a potential explanation of its higher expression in CML. Based on these results, we expect the TD module of Pindel will play an important role in data analysis of cancer genomic sequencing data.

**Figure 3.**
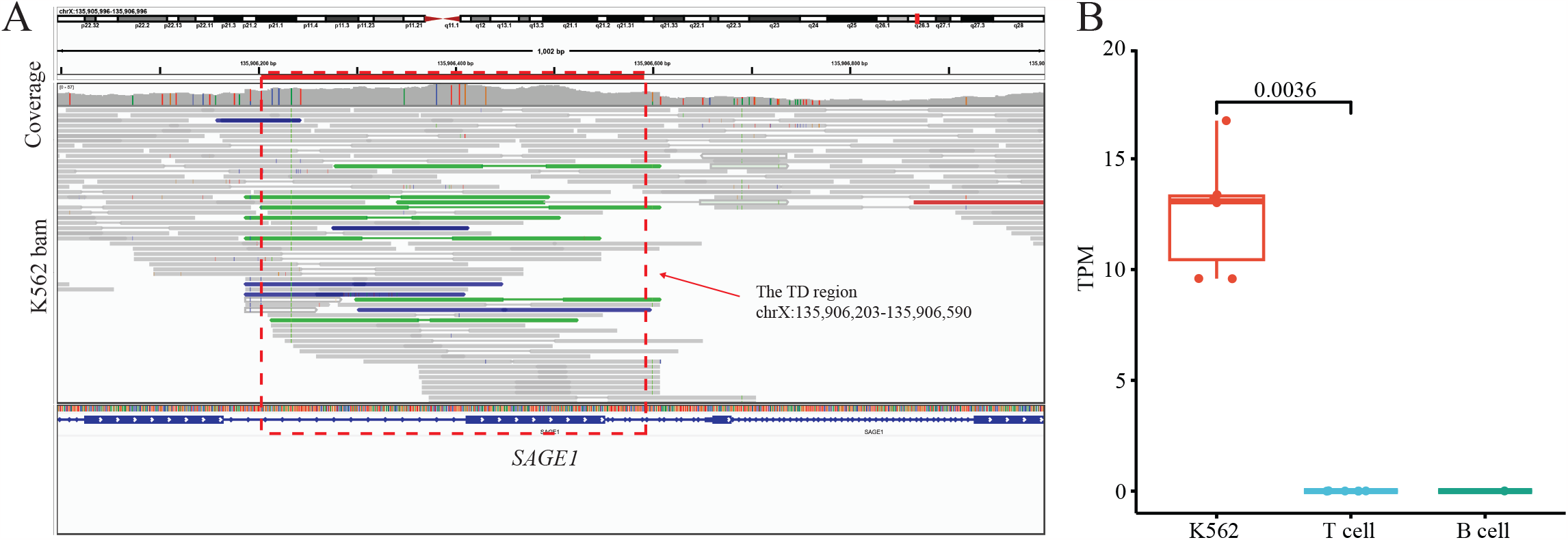
SAGE1 related TD detected from K562 cell line. **A**. The IGV screenshot of TD located at chrX:135,906,203-135,906,590 of K562 cell line. **B**. The gene expression of *SAGE1* in K562, T cell, and B cell. The *p*-value is calculated by Wilcoxon Rank Sum and Signed Rank test. We cannot calculate the *p*-value between the expression of K562 and B cell due to insufficient number of data points (only one RNA-seq data of B cell acquired from ENCODE). TPM: transcript per million.

## Discussion

In this study, we developed a TD detection model as Pindel-TD by specifically optimizing the pattern growth approach in Pindel. We redesigned the search strategies of the minimum and maximum unique substring for different sized TDs, resulting in the high and robust performance of TD detection on a wide size range. We compared Pindel-TD with widely-used SV detection methods, including Manta, Delly, and Lumpy, on simulation and HG002 sequencing data, demonstrating the outperformance of Pindel-TD. We further applied Pindel-TD on sequencing data of K562 cell line, and detected a TD containing the seventh exon of *SAGE1*, potentially explained its high expression, demonstrating the potential application of Pindel-TD on cancer studies.

We evaluated the false discovery rate based on the simulation data (Table S1). However, it is hard to evaluate the false positive rate due to lacking of negative TDs for either simulation or real data. Split-reads were used to improve the break point resolution. However, such split-reads only account for a small part of the total number of reads that related with TDs, limiting the genotype analysis.

Currently, Pindel is developed for short-read sequencing data while does not support the data from long-read sequencing, which has been selected as the Method of Year 2022 [32]. Although long read sequencing powers genomic studies, *e*.*g*., genome assembly [33-36] and structural variation detection [37, 38], its high cost limits the accessibility to more research communities, *e*.*g*., large cohort and clinicogenomics studies. Therefore, methods developed for short-read sequencing data still has its unreplaceable values for genomic community. In addition, we are trying to develop the new version of Pindel to support long-read and short-read sequencing data for detecting both simple and complex SVs.

## Data availability

The GRCh38 was downloaded from https://ftp.1000genomes.ebi.ac.uk/vol1/ftp/technical/reference/GRCh38_reference_genome/GRCh38_full_analysis_set_plus_decoy_hla.fa. The pair-end sequencing data of the widely used HG002 cell line was downloaded from https://ftp-trace.ncbi.nlm.nih.gov/ReferenceSamples/giab/data/AshkenazimTrio/HG002_NA24385_son/NIST_Illumina_2x250bps/reads/. The v0.6 SV benchmark set for HG002 is downloaded from https://ftp-trace.ncbi.nlm.nih.gov/ReferenceSamples/giab/data/AshkenazimTrio/analysis/NIST_SVs_Integration_v0.6/HG002_SVs_Tier1_v0.6.vcf.gz. The HiFi long read sequencing data of HG002 was downloaded from https://www.ncbi.nlm.nih.gov/sra/ under Accessions: SRR18239004, SRR18239005, SRR18239006, and SRR18239007. The repetitive regions of GRCh37 were downloaded from UCSC table browser (https://genome.ucsc.edu/cgi-bin/hgTables) with tracks of Simple Repeats and RepeatMasker. The short-read sequencing data of K562 cancer cell line was downloaded from ENCODE (https://www.encodeproject.org/experiments/ENCSR045NDZ/). The gene expression of K562, T cell, and B cell were downloaded from ENCODE (https://www.encodeproject.org/) under Accessions: ENCFF718ZTE, ENCFF659CXW, ENCFF922MEO, ENCFF922MEO, ENCFF006PZD, and ENCFF047KBI for K562, ENCFF637STR, ENCFF924TOQ, ENCFF492RBW, ENCFF090XWE, ENCFF192WQE, and ENCFF787FKA for T cell, and ENCFF192WQE for B cell.

## Code availability

Pindel-TD is available at https://github.com/xjtu-omics/pindel. It is free for non-commercial use by academic, government, and non-profit/not-for-profit institutions. A commercial version of the software is available and licensed through Xi’an Jiaotong University. For more information, please contact.

## Author’s contributions

KY conceived of, designed, and supervised the study XY, GZ, KY developed the pattern growth approach and implemented the source code XY, GZ, PJ and SW evaluated the performances of Pindel-TD. XY, GZ, and KY wrote the manuscript. All authors contributed to critical revision of the manuscript, read, and approved the final version.

## Acknowledgments

This work was supported by the National Key R&D Program of China (2022YFC3400300), the National Natural Science Foundation of China (62172325, 32125009, 32070663), the Key Construction Program of the National ‘985′ Project, and the Fundamental Research Funds for the Central Universities.

## Supplementary tables

Table S1. The simulated TDs.

Table S2. The benchmarked TDs in HG002.

Table S3. The performance evaluation of SV methods on simulated TDs. Table S4. The 1,162 TDs detected by Pindel-TD from HG002.

Table S5. The precision of detecting TDs on HG002 sequencing data.

Table S6. The recall of different methods on HG002 benchmarked TDs for different TR content.

Table S7. The TDs detected from K562 cell line.

Table S8. Gene expression of *SAGE1in* K562 cell line, T cell and B cell from ENCODE.

## Supplementary figures

See in supplementary materials

## References

[1] Ho SS, Urban AE, Mills RE. Structural variation in the sequencing era. Nat Rev Genet 2020 21:171–89.

[2] Alkan C, Coe BP, Eichler EE. Genome structural variation discovery and genotyping. Nat Rev Genet 2011 12:363–76.

[3] Rogers RL, Shao L, Thornton KR. Tandem duplications lead to novel expression patterns through exon shuffling in Drosophila yakuba. PLoS Genet 2017 13:e1006795.

[4] Xu Z, Pu X, Gao R, Demurtas OC, Fleck SJ, Richter M, et al. Tandem gene duplications drive divergent evolution of caffeine and crocin biosynthetic pathways in plants. BMC Biol 2020 18:63.

[5] Miller DE, Squire A, Bennett JT. A child with autism, behavioral issues, and dysmorphic features found to have a tandem duplication within CTNND2 by mate-pair sequencing. Am J Med Genet A 2020 182:543–7.

[6] Menghi F, Inaki K, Woo X, Kumar PA, Grzeda KR, Malhotra A, et al. The tandem duplicator phenotype as a distinct genomic configuration in cancer. Proc Natl Acad Sci U S A 2016 113:E2373–82.

[7] Menghi F, Barthel FP, Yadav V, Tang M, Ji B, Tang Z, et al. The Tandem Duplicator Phenotype Is a Prevalent Genome-Wide Cancer Configuration Driven by Distinct Gene Mutations. Cancer Cell 2018 34:197–210 e5.

[8] Ye K, Schulz MH, Long Q, Apweiler R, Ning Z. Pindel: a pattern growth approach to detect break points of large deletions and medium sized insertions from paired-end short reads. Bioinformatics 2009 25:2865–71.

[9] Ye K, Wang J, Jayasinghe R, Lameijer EW, McMichael JF, Ning J, et al. Systematic discovery of complex insertions and deletions in human cancers. Nat Med 2016 22:97–104.

[10] Rausch T, Zichner T, Schlattl A, Stutz AM, Benes V, Korbel JO. DELLY: structural variant discovery by integrated paired-end and split-read analysis. Bioinformatics 2012 28:i333–i9.

[11] Layer RM, Chiang C, Quinlan AR, Hall IM. LUMPY: a probabilistic framework for structural variant discovery. Genome Biol 2014 15:R84.

[12] Lee JJ, Park S, Park H, Kim S, Lee J, Lee J, et al. Tracing Oncogene Rearrangements in the Mutational History of Lung Adenocarcinoma. Cell 2019 177:1842–57 e21.

[13] Lin J, Yang X, Kosters W, Xu T, Jia Y, Wang S, et al. Mako: A Graph-based Pattern Growth Approach to Detect Complex Structural Variants. Genomics Proteomics Bioinformatics 2022 20:205–18.

[14] Lin J, Wang S, Audano PA, Meng D, Flores JI, Kosters W, et al. SVision: a deep learning approach to resolve complex structural variants. Nat Methods 2022 19:1230–3.

[15] Xi R, Kim TM, Park PJ. Detecting structural variations in the human genome using next generation sequencing. Brief Funct Genomics 2010 9:405–15.

[16] Liu Y, Huang Y, Wang G, Wang Y. A deep learning approach for filtering structural variants in short read sequencing data. Brief Bioinform 2021 22.

[17] Zook JM, Hansen NF, Olson ND, Chapman L, Mullikin JC, Xiao C, et al. A robust benchmark for detection of germline large deletions and insertions. Nat Biotechnol 2020 38:1347–55.

[18] Chen X, Schulz-Trieglaff O, Shaw R, Barnes B, Schlesinger F, Kallberg M, et al. Manta: rapid detection of structural variants and indels for germline and cancer sequencing applications. Bioinformatics 2016 32:1220–2.

[19] Dong J, Qi M, Wang S, Yuan X. DINTD: Detection and Inference of Tandem Duplications From Short Sequencing Reads. Front Genet 2020 11:924.

[20] Consortium EP, Moore JE, Purcaro MJ, Pratt HE, Epstein CB, Shoresh N, et al. Expanded encyclopaedias of DNA elements in the human and mouse genomes. Nature 2020 583:699–710.

[21] Sosic M, Sikic M. Edlib: a C/C ++ library for fast, exact sequence alignment using edit distance. Bioinformatics 2017 33:1394–5.

[22] Bolognini D, Sanders A, Korbel JO, Magi A, Benes V, Rausch T. VISOR: a versatile haplotype-aware structural variant simulator for short- and long-read sequencing. Bioinformatics 2020 36:1267–9.

[23] Li H, Durbin R. Fast and accurate short read alignment with Burrows-Wheeler transform. Bioinformatics 2009 25:1754–60.

[24] Krumsiek J, Arnold R, Rattei T. Gepard: a rapid and sensitive tool for creating dotplots on genome scale. Bioinformatics 2007 23:1026–8.

[25] Sedlazeck FJ, Rescheneder P, Smolka M, Fang H, Nattestad M, von Haeseler A, et al. Accurate detection of complex structural variations using single-molecule sequencing. Nat Methods 2018 15:461–8.

[26] English AC, Menon VK, Gibbs RA, Metcalf GA, Sedlazeck FJ. Truvari: refined structural variant comparison preserves allelic diversity. Genome Biol 2022 23:271.

[27] Benson G. Tandem repeats finder: a program to analyze DNA sequences. Nucleic Acids Res 1999 27:573–80.

[28] Zhang Y, Yu X, Liu Q, Gong H, Chen AA, Zheng H, et al. SAGE1: a Potential Target Antigen for Lung Cancer T-Cell Immunotherapy. Mol Cancer Ther 2021 20:2302–13.

[29] Deniz O, Ahmed M, Todd CD, Rio-Machin A, Dawson MA, Branco MR. Endogenous retroviruses are a source of enhancers with oncogenic potential in acute myeloid leukaemia. Nat Commun 2020 11:3506.

[30] Dufva O, Polonen P, Bruck O, Keranen MAI, Klievink J, Mehtonen J, et al. Immunogenomic Landscape of Hematological Malignancies. Cancer Cell 2020 38:380–99 e13.

[31] Wang D, Yang H, Zhang Y, Hu R, Hu D, Wang Q, et al. Inhibition of cystathionine beta-synthase promotes apoptosis and reduces cell proliferation in chronic myeloid leukemia. Signal Transduct Target Ther 2021 6:52.

[32] Method of the Year 2022: long-read sequencing. Nat Methods 2023 20:1.

[33] Wang B, Yang X, Jia Y, Xu Y, Jia P, Dang N, et al. High-quality Arabidopsis thaliana Genome Assembly with Nanopore and HiFi Long Reads. Genomics Proteomics Bioinformatics 2022 20:4–13.

[34] Aganezov S, Yan SM, Soto DC, Kirsche M, Zarate S, Avdeyev P, et al. A complete reference genome improves analysis of human genetic variation. Science 2022 376:eabl3533.

[35] Yang X, Gao S, Guo L, Wang B, Jia Y, Zhou J, et al. Three chromosome-scale Papaver genomes reveal punctuated patchwork evolution of the morphinan and noscapine biosynthesis pathway. Nat Commun 2021 12:6030.

[36] Guo L, Winzer T, Yang X, Li Y, Ning Z, He Z, et al. The opium poppy genome and morphinan production. Science 2018 362:343–7.

[37] Audano PA, Sulovari A, Graves-Lindsay TA, Cantsilieris S, Sorensen M, Welch AE, et al. Characterizing the Major Structural Variant Alleles of the Human Genome. Cell 2019 176:663–75 e19.

[38] Ebert P, Audano PA, Zhu Q, Rodriguez-Martin B, Porubsky D, Bonder MJ, et al. Haplotype-resolved diverse human genomes and integrated analysis of structural variation. Science 2021 372.

