## Supplementary figures and images for "Pindel-TD: a tandem duplication detector based on a pattern growth approach"

### Supplementary Figure 1

#### Supplementary Figure

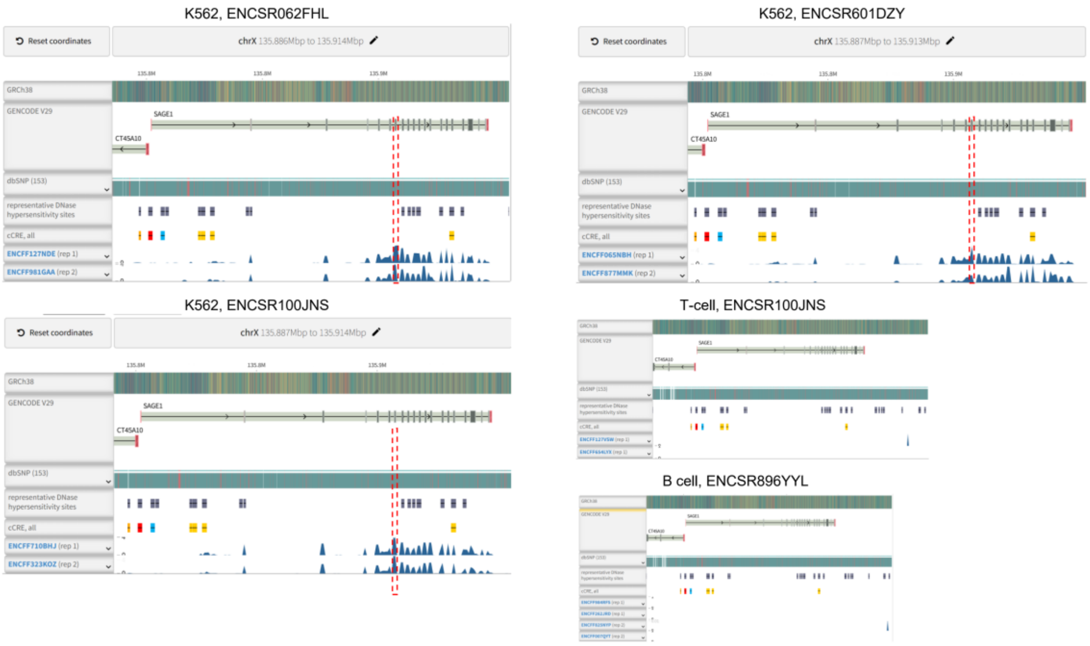


**Fig. S1. RNA-seq coverage of *SAGE1* in K562 cell line, T cell and B cell from ENCODE.**
